## Supplementary Material for "ACMTF-R: supervised multi-omics data integration uncovering shared and distinct outcome-associated variation"

**Supplementary Methods**

*Gradient function of ACMTF*

In the case where the first mode is shared across all data blocks, and assuming that every term in $f\left( \alpha\mathbf{,}\beta\mathbf{,}\varepsilon\boldsymbol{,\Lambda,}\boldsymbol{A},\boldsymbol{B}^{\left( 1 \right)},\boldsymbol{C}^{\left( 1 \right)},\ldots,\boldsymbol{B}^{\left( P \right)},\boldsymbol{C}^{\left( P \right)} \right)$ is multiplied by $1/2$, the gradient of the loss function $\nabla f$ with size $F\left( I+J^{(1)}+K^{(1)}+\ldots+J^{\left( P \right)}+K^{(P)}+P \right)\times1$ becomes

$$\begin{aligned} \nabla f=\left[ \text{vec}\left( \frac{\partial f}{\partial\boldsymbol{A}} \right)^{T}\text{vec}\left( \frac{\partial f}{\partial\boldsymbol{B}^{\left( 1 \right)}} \right)^{T}\text{vec}\left( \frac{\partial f}{\partial\boldsymbol{C}^{\left( 1 \right)}} \right)^{T}\ldots\text{vec}\left( \frac{\partial f}{\partial\boldsymbol{B}^{\left( P \right)}} \right)^{T}\text{vec}\left( \frac{\partial f}{\partial\boldsymbol{C}^{\left( P \right)}} \right)^{T}{\frac{\partial f}{\partial\boldsymbol{\lambda}_{1}}}^{T}{\frac{\partial f}{\partial\boldsymbol{\lambda}_{2}}}^{T}\ldots{\frac{\partial f}{\partial\boldsymbol{\lambda}_{P}}}^{T} \right]^{T}\#\left( 5 \right) \end{aligned}$$

where

$$\frac{\partial f}{\partial\boldsymbol{A}}=\sum_{p=1}^{P} ({\hat{\boldsymbol{X}}}_{a}^{\left( p \right)}-\boldsymbol{X}_{a}^{(p)})(\boldsymbol{\lambda}_{p}^{T}⨀\boldsymbol{C}^{\left( p \right)}\boldsymbol{⨀}\boldsymbol{B}^{\left( p \right)}\boldsymbol{)}+\alpha\left( \boldsymbol{A}-\bar{\boldsymbol{A}} \right),$$

$$\frac{\partial f}{\partial\boldsymbol{B}^{(p)}}=\sum_{p=1}^{P} ({\hat{\boldsymbol{X}}}_{b}^{\left( p \right)}-\boldsymbol{X}_{b}^{(p)})(\boldsymbol{\lambda}_{p}^{T}⨀\boldsymbol{C}^{\left( p \right)}\boldsymbol{⨀A)}+\alpha\left( \boldsymbol{B}^{(p\boldsymbol{)}}-{\bar{\boldsymbol{B}}}^{(p)} \right),$$

$$\frac{\partial f}{\partial\boldsymbol{C}^{(p)}}=\sum_{p=1}^{P} ({\hat{\boldsymbol{X}}}_{c}^{\left( p \right)}-\boldsymbol{X}_{c}^{(p)})(\boldsymbol{\lambda}_{p}^{T}⨀\boldsymbol{B}^{\left( p \right)}\boldsymbol{⨀A)}+\alpha\left( \boldsymbol{C}^{(p\boldsymbol{)}}-{\bar{\boldsymbol{C}}}^{(p)} \right),$$

$$\frac{\partial f}{\partial\lambda_{pf}}=\left( {\hat{\underline{\boldsymbol{X}}}}^{(p)}-{\underline{\boldsymbol{X}}}^{(p)} \right)\times_{1}\boldsymbol{a}_{f}\times_{2}\boldsymbol{b}_{f}^{(p)}\times_{3}\boldsymbol{c}_{f}^{(p)}+\frac{\beta}{2}\frac{\lambda_{pf}}{\sqrt{\lambda_{pf}^{2}+\epsilon}},$$

and $\times_{n}$is the tensor-vector product in the $n$-th mode as defined in [35]. $\bar{\boldsymbol{A}}$ corresponds to the matrix $\boldsymbol{A}$ with columns divided by their 2-norms. The derivation of this gradient is reported in [18]. The acmtf_opt() function runs this model in the provided R package.

*Gradient function of ACMTF-R*

Assuming that the first mode is shared across all data blocks and that every term in $f\left( \alpha\mathbf{,}\beta\mathbf{,}\varepsilon\mathbf{,}\pi\boldsymbol{,\Lambda,}\boldsymbol{A},\boldsymbol{B}^{\left( 1 \right)},\boldsymbol{C}^{\left( 1 \right)},\ldots,\boldsymbol{B}^{\left( P \right)},\boldsymbol{C}^{\left( P \right)} \right)$ is multiplied by $1/2$, the gradient of the loss function becomes

$$\begin{aligned} \nabla f=\left[ \text{vec}\left( \frac{\partial f}{\partial\boldsymbol{A}} \right)^{T}\text{vec}\left( \frac{\partial f}{\partial\boldsymbol{B}^{\left( 1 \right)}} \right)^{T}\text{ vec}\left( \frac{\partial f}{\partial\boldsymbol{C}^{\left( 1 \right)}} \right)^{T}\ldots\text{vec}\left( \frac{\partial f}{\partial\boldsymbol{B}^{\left( P \right)}} \right)^{T}\text{vec}\left( \frac{\partial f}{\partial\boldsymbol{C}^{\left( P \right)}} \right)^{T}{\frac{\partial f}{\partial\boldsymbol{\lambda}_{1}}}^{T}{\frac{\partial f}{\partial\boldsymbol{\lambda}_{2}}}^{T}\ldots{\frac{\partial f}{\partial\boldsymbol{\lambda}_{P}}}^{T} \right]^{T}\#\left( 8 \right) \end{aligned}$$

where

$$\frac{\partial f}{\partial\boldsymbol{A}}=\pi\sum_{p=1}^{P} ({\hat{\boldsymbol{X}}}_{a}^{\left( p \right)}-\boldsymbol{X}_{a}^{(p)})\left( \boldsymbol{\lambda}_{\boldsymbol{p}}^{T}⨀\boldsymbol{C}^{\left( p \right)}\boldsymbol{⨀}\boldsymbol{B}^{\left( p \right)} \right)+\left( 1-\pi\right)\left( \boldsymbol{A\rho-y} \right)\boldsymbol{\rho}^{T}\boldsymbol{+}\alpha\left( \boldsymbol{A}-\bar{\boldsymbol{A}} \right),$$

$$\frac{\partial f}{\partial\boldsymbol{B}^{\boldsymbol{(p)}}}=\pi\sum_{p=1}^{P} ({\hat{\boldsymbol{X}}}_{b}^{\left( p \right)}-\boldsymbol{X}_{b}^{(p)})(\boldsymbol{\lambda}_{p}^{T}⨀\boldsymbol{C}^{\left( p \right)}\boldsymbol{⨀A)}+\alpha\left( \boldsymbol{B}^{(p\boldsymbol{)}}-{\bar{\boldsymbol{B}}}^{(p)} \right),$$

$$\frac{\partial f}{\partial\boldsymbol{C}^{\boldsymbol{(p)}}}=\pi\sum_{p=1}^{P} ({\hat{\boldsymbol{X}}}_{c}^{\left( p \right)}-\boldsymbol{X}_{c}^{(p)})(\boldsymbol{\lambda}_{p}^{T}⨀\boldsymbol{B}^{\left( p \right)}\boldsymbol{⨀A)}+\alpha\left( \boldsymbol{C}^{(p\boldsymbol{)}}-{\bar{\boldsymbol{C}}}^{(p)} \right),$$

$$\frac{\partial f}{\partial\lambda_{pf}}=\left( {\hat{\underline{\boldsymbol{X}}}}^{(p)}-{\underline{\boldsymbol{X}}}^{(p)} \right)\times_{1}\boldsymbol{a}_{f}\times_{2}\boldsymbol{b}_{f}^{(p)}\times_{3}\boldsymbol{c}_{f}^{(p)}+\frac{\beta}{2}\frac{\lambda_{pf}}{\sqrt{\lambda_{pf}^{2}+\epsilon}},$$

and $\times_{n}$ is the tensor-vector product in the n-th mode as defined in [35]. $\bar{\boldsymbol{A}}$ corresponds to the matrix $\boldsymbol{A}$ with the columns divided by their 2-norms. The derivation of this gradient is reported in **Appendix 1**. The acmtfr_opt() function runs this algorithm in the provided R package.

*Missing data*

Missing data is an unfortunate reality of sampling biological data, especially for microbiome data. In cohort studies, some participants might not turn up for sampling at one or more time points, or they may drop out altogether. Low library sizes may cause some samples to be removed from the data. Such missing data ends up as rows of missing data in the three-way array. Fortunately, the ACMTF and ACMTF-R function can be adapted to accommodate this. This section highlights the mathematical adaptations required for ACMTF and ACMTF-R to work appropriately in the presence of missing data. First, the definitions of the binary weight array $\boldsymbol{W}$ and the weighted norm $\left\| \cdot\right\|_{W}$ are given. Then the modifications for ACMTF and ACMTF-R are outlined.

*Binary weight array W*

All of the mathematical adaptations presented in this section required a binary three-way array of weights $\underline{\boldsymbol{W}}$ encoding the non-missing values in the data $\underline{\boldsymbol{X}}$ where

$$\begin{aligned} w_{ijk}=\left\{ \begin{aligned} 0 \text{if }x_{ijk}\text{is missing} \\ 1 \text{otherwise,} \end{aligned} \right. \#\left( 1 \right) \end{aligned}$$

and the array element $x_{ijk}$ in $\underline{\boldsymbol{X}}$ corresponding to subject $i$ and feature $j$ at time point $k$ is used to determine the corresponding array element $w_{ijk}$ in $\underline{\boldsymbol{W}}$ and a two-way array of weights is defined equivalently [1, 2].

*Weighted norm*

The weighted norm of a three-way array $\underline{\boldsymbol{X}}$ is defined as

$$\begin{aligned} \left\| \underline{\boldsymbol{X}} \right\|_{W}=\left\| \underline{\boldsymbol{W}}*\underline{\boldsymbol{X}} \right\| \#\left( 2 \right) \end{aligned}$$

where the weight array $\underline{\boldsymbol{W}}$ is used to encode the non-missing array elements in $\underline{\boldsymbol{X}}$ [1, 2]. An equivalent definition is used for the two-way and vector cases.

*Weighted ACMTF*

In ACMTF, the weighted calculation of the loss only affects the first term

$$\begin{aligned} f\left( \alpha\mathbf{,}\beta\mathbf{,}\varepsilon\boldsymbol{,\Lambda,}\boldsymbol{A},\boldsymbol{B}^{\left( 1 \right)},\boldsymbol{C}^{\left( 1 \right)},\ldots,\boldsymbol{B}^{\left( P \right)},\boldsymbol{C}^{\left( P \right)} \right)\mathbf{=}\sum_{p=1}^{P} \left\| {\underline{\boldsymbol{X}}}^{\left( p \right)}-{\underline{\hat{\boldsymbol{X}}}}^{\left( p \right)} \right\|_{W}^{2} \\ + \beta\sum_{p=1}^{P} \sum_{f=1}^{F} \sqrt{\lambda_{fp}^{2}+\epsilon} \\ + \alpha\sum_{p=1}^{P} \sum_{f=1}^{F} \left( \left\| \boldsymbol{a}_{f} \right\|-1 \right)^{2} \\ + \alpha\sum_{p=1}^{P} \sum_{f=1}^{F} \left( \left\| \boldsymbol{b}_{f}^{(p)} \right\|-1 \right)^{2} \\ + \alpha\sum_{p=1}^{P} \sum_{f=1}^{F} \left( \left\| \boldsymbol{c}_{f}^{(p)} \right\|-1 \right)^{2} \#\left( 3 \right) \end{aligned}$$

and the gradient function is unaffected [3].

*Weighted ACMTF-R*

In ACMTF-R, the weighted calculation of the loss only affects the first term

$$\begin{aligned} f\left( \alpha\mathbf{,}\beta\mathbf{,}\varepsilon\mathbf{,}\pi\boldsymbol{,\Lambda,}\boldsymbol{A},\boldsymbol{B}^{\left( 1 \right)},\boldsymbol{C}^{\left( 1 \right)},\ldots,\boldsymbol{B}^{\left( P \right)},\boldsymbol{C}^{\left( P \right)} \right)\mathbf{=}\pi\sum_{p=1}^{P} \left\| {\underline{\boldsymbol{X}}}^{\left( p \right)}-{\underline{\hat{\boldsymbol{X}}}}^{\left( p \right)} \right\|_{W}^{2} \\ + \left( 1-\pi\right)\left\| \boldsymbol{y-A\rho} \right\|^{2} \\ + \beta\sum_{p=1}^{P} \sum_{f=1}^{F} \sqrt{\lambda_{fp}^{2}+\epsilon} \\ + \alpha\sum_{p=1}^{P} \sum_{f=1}^{F} \left( \left\| \boldsymbol{a}_{f} \right\|-1 \right)^{2} \\ + \alpha\sum_{p=1}^{P} \sum_{f=1}^{F} \left( \left\| \boldsymbol{b}_{f}^{(p)} \right\|-1 \right)^{2} \\ + \alpha\sum_{p=1}^{P} \sum_{f=1}^{F} \left( \left\| \boldsymbol{c}_{f}^{(p)} \right\|-1 \right)^{2} \#\left( 4 \right) \end{aligned} \left( 6 \right)$$

and the gradient function is unaffected (**Appendix 1**).

When $y_{new}$ is calculated for a new sample $\boldsymbol{x}_{new}$ containing some missing values, the weighted calculation of the joint scores becomes

$$\boldsymbol{a}_{\boldsymbol{new}}\boldsymbol{=}\boldsymbol{W}\boldsymbol{*}\boldsymbol{Z}^{\boldsymbol{+}}\text{vec(}x_{new})$$

where $\boldsymbol{Z}^{\boldsymbol{+}}$ is the Moore-Penrose inverse of $\boldsymbol{Z}$ and $\boldsymbol{W}$ is a binary weight matrix encoding the non-missing data for $\text{vec(}x_{new})$. Subsequently $y_{new}$ is found by using the regression coefficients $\boldsymbol{\rho}$ as in the non-weighted version of the procedure.

*Model evaluation - Fit*

Model evaluation is typically done using the percentage of variance explained. In the context of ACMTF-R this can be done for a data block ${\underline{\boldsymbol{X}}}^{(p)}$ and $\boldsymbol{y}$ through

$$\begin{aligned} Fit_{{\underline{\boldsymbol{X}}}^{(p)}}\left( \% \right)=\left( 1-\frac{\left\| {\underline{\boldsymbol{X}}}^{\left( p \right)}-{\underline{\hat{\boldsymbol{X}}}}^{\left( p \right)} \right\|^{2}}{\left\| {\underline{\boldsymbol{X}}}^{(p)} \right\|} \right)*100 \#\left( 10 \right) \end{aligned}$$

and

$$\begin{aligned} Fit_{\boldsymbol{y}}\left( \% \right)=\left( 1-\frac{\left\| \boldsymbol{y}-\hat{\boldsymbol{y}} \right\|^{2}}{\left\| \boldsymbol{y} \right\|^{2}} \right)*100. \#\left( 11 \right) \end{aligned}$$

While variance explained gives an overall impression of the model fit, it does not give a good indication of whether the input loadings have been found, nor if the correct CLD structure has been found, especially in the context of noise.

When missing data is present, the fit can be computed for a data block ${\underline{\boldsymbol{X}}}^{(p)}$ through

$$\begin{aligned} Fit_{{\underline{\boldsymbol{X}}}^{(p)}}\left( \% \right)=\left( 1-\frac{\left\| \boldsymbol{X}^{\left( p \right)}-{\hat{\boldsymbol{X}}}^{\left( p \right)} \right\|_{W}^{2}}{\left\| \boldsymbol{X}^{\left( p \right)} \right\|_{W}^{2}} \right)*100 \#\left( 10 \right) \end{aligned}$$

and the calculation of the variation explained in $\boldsymbol{y}$ remains unchanged.

*Model evaluation – Lambda Similarity Index*

We define the Lambda Similarity Index ($LSI$) to compare the found lambda matrix $\hat{\boldsymbol{\Lambda}}$ to the true lambda matrix $\boldsymbol{\Lambda}$ using an inverted sum-of-squared residuals approach

$$\begin{aligned} LSI=\frac{1}{1+\sum_{p=1}^{P} \sum_{f=1}^{F} \left( \lambda_{pf}-\hat{\lambda}_{pf} \right)^{2}} \#\left( 13 \right) \end{aligned}$$

where $p$ is the block index and $f$ is the component number. This metric has the range (0,$1$], where $LSI=1$ corresponds to the ACMTF-R model finding the correct CLD-structure and $LSI\to0$ corresponds to the ACMTF-R model not finding the correct CLD-structure in any way. LSI does not take the recovered factor loadings into account.

*Model evaluation – Matching components*

Both FMS and LSI require a best matching of components due to the permutational freedom of the model. However, the chosen permutation should be the same for both metrics. This is done by first computing FMS and LSI for all pairwise combinations of factors and subsequently using the Hungarian algorithm [4, 5] to find the matching of components that maximises both metrics. The combination of FMS and LSI will be used to evaluate CLD-structure and factor recovery of ACMTF-R for all simulations.

**Supplementary Figures**

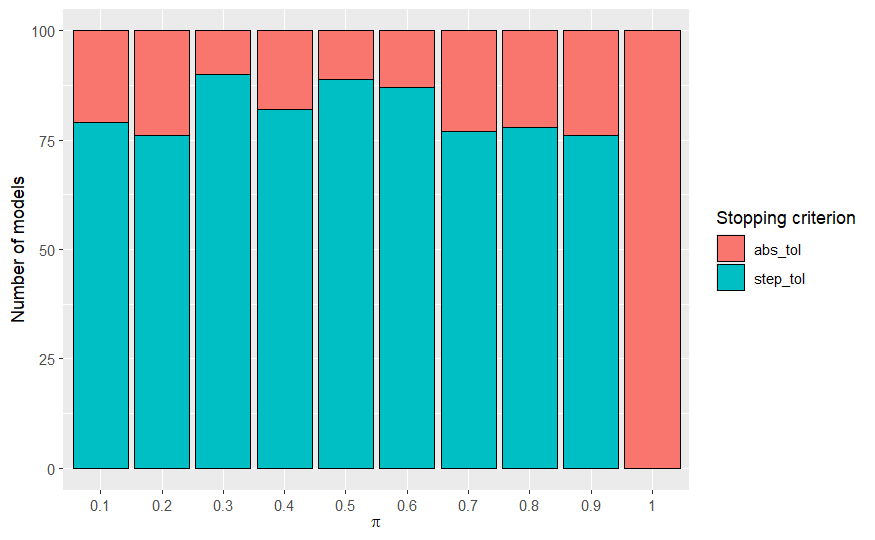

**Supplementary Figure 1**. Overview of the stopping criterion that terminated the ACMTF-R line search algorithm in the first simulation. abs_tol: minimum value of the loss function. step_tol: absolute tolerance for the size of the parameter update.

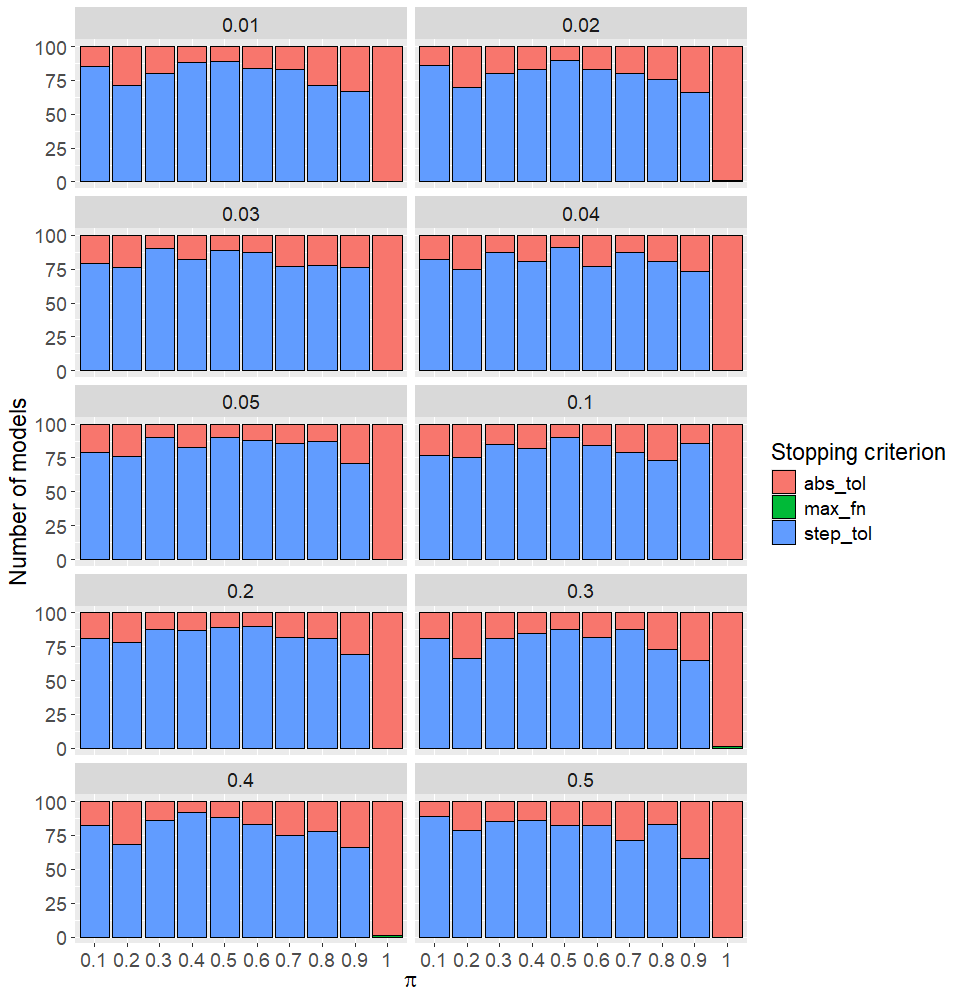
 **Supplementary Figure 2**. Overview of the stopping criterion that terminated the ACMTF-R line search algorithm in the second simulation. Plots are faceted by the size of the hidden component ($\delta$). abs_tol: minimum value of the loss function. step_tol: absolute tolerance for the size of the parameter update. max_fn: maximum number of function evaluations.

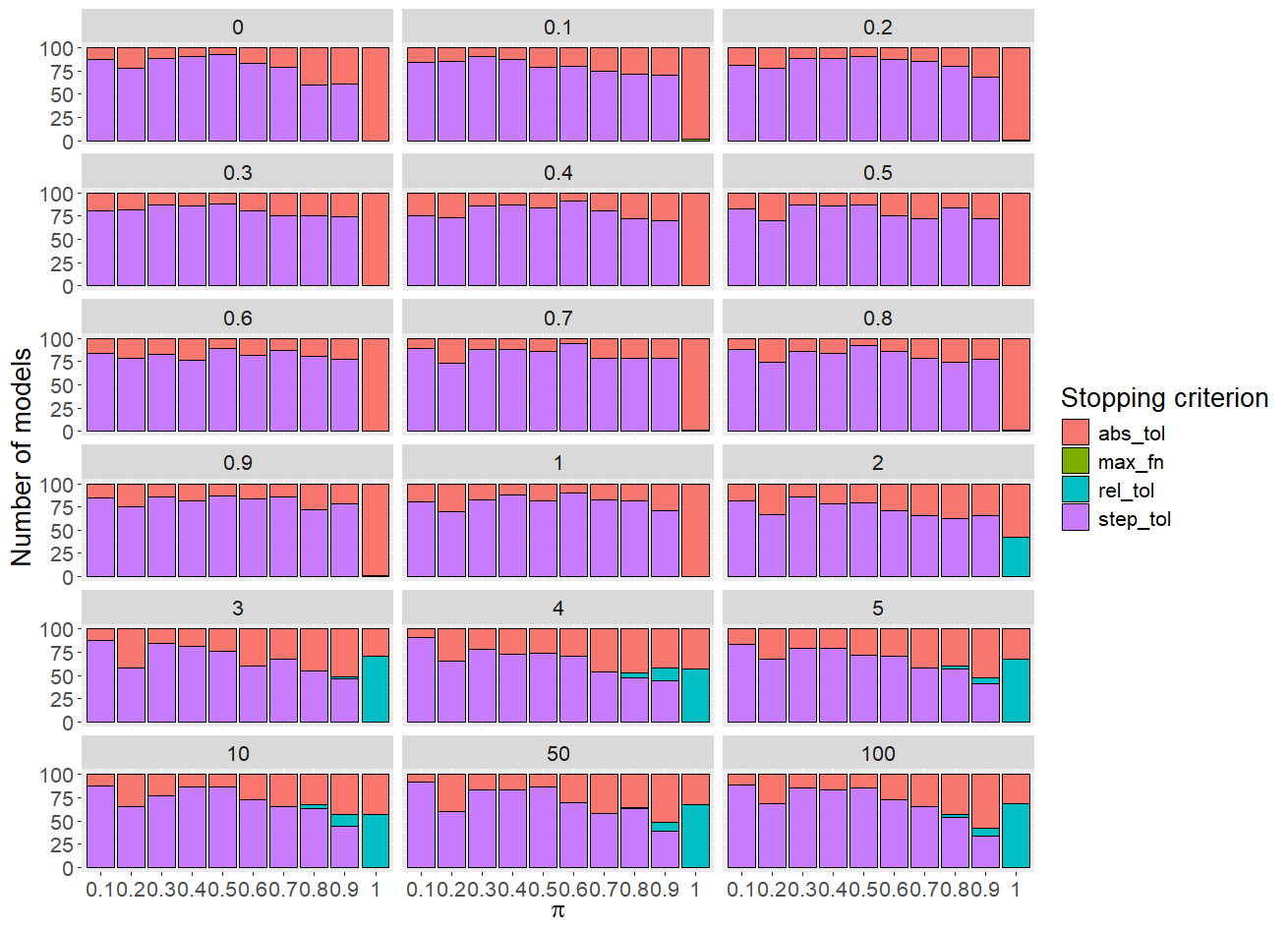

**Supplementary Figure 3.** Overview of the stopping criterion that terminated the ACMTF-R line search algorithm in the third simulation. Plots are faceted by the noise level on X ($\eta_{X}$). abs_tol: minimum value of the loss function. rel_tol: relative change of the loss function. step_tol: absolute tolerance for the size of the parameter update. max_fn: maximum number of function evaluations.

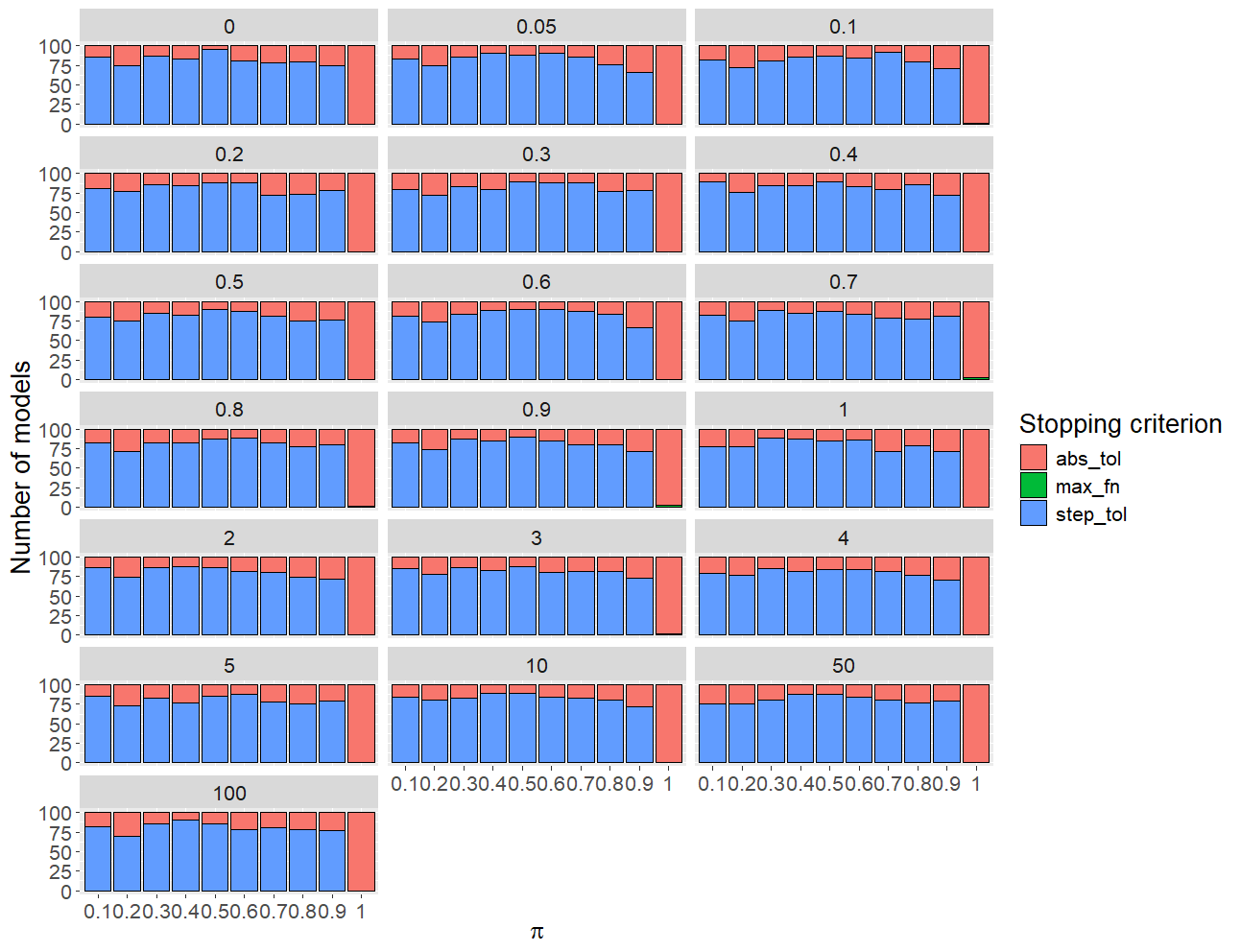

**Supplementary Figure 4.** Overview of the stopping criterion that terminated the ACMTF-R line search algorithm in the fourth simulation. Plots are faceted by the noise level on $\boldsymbol{y}$ ($\eta_{y}$). abs_tol: minimum value of the loss function. step_tol: absolute tolerance for the size of the parameter update. max_fn: maximum number of function evaluations.

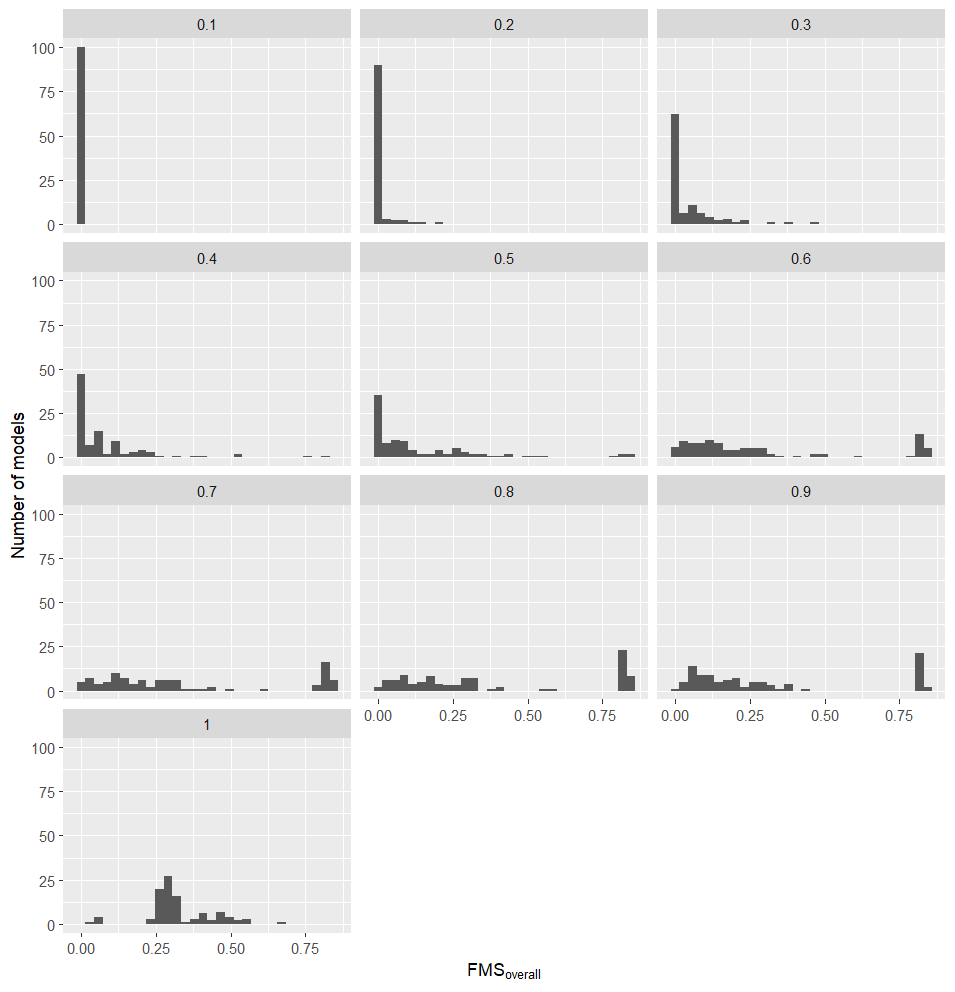

**Supplementary Figure 5.** Histogram of the $FMS_{overall}$ observed across the randomly initialised ACMTF and ACMTF-R models in the first simulation. Plots are faceted by $\pi$ value and each contain 100 randomly initialised models. ACMTF corresponds to $\pi=1$.

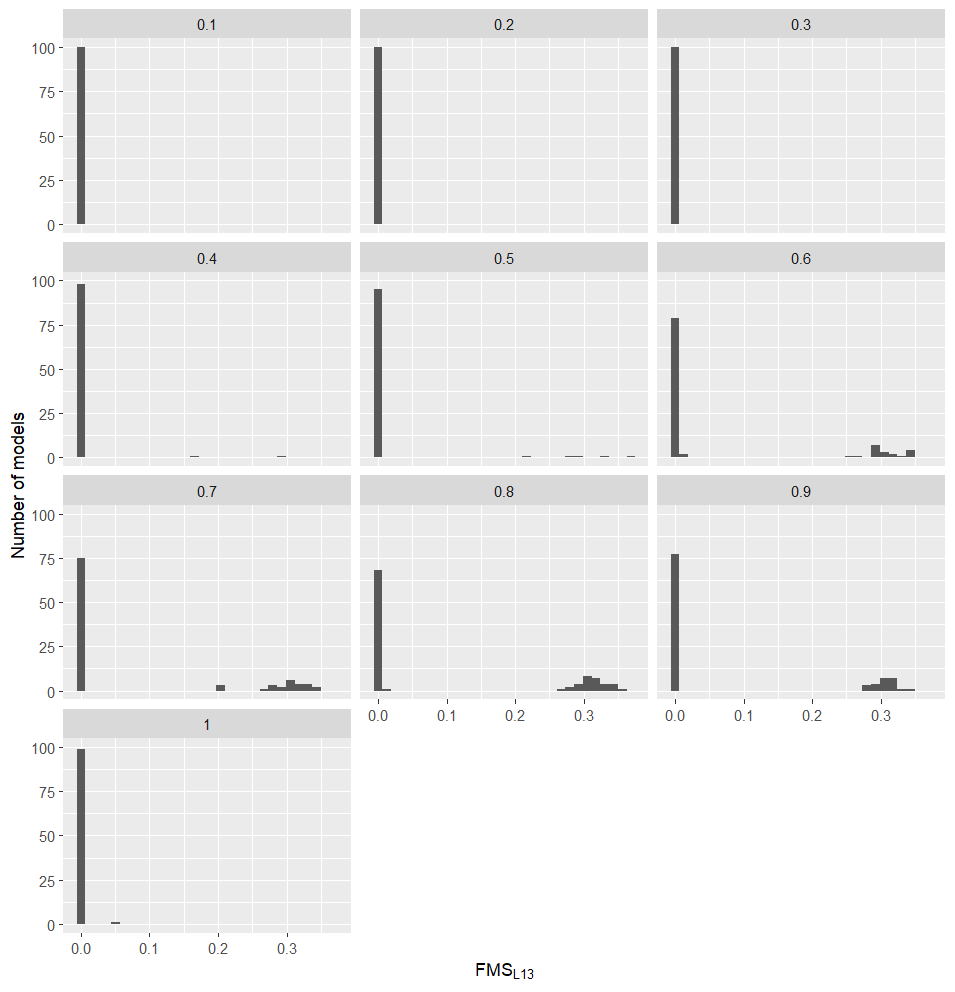

**Supplementary Figure 6.** Histogram of the $FMS_{L13}$ observed across the randomly initialised ACMTF and ACMTF-R models in the first simulation. Plots are faceted by $\pi$ value and each contain 100 randomly initialised models. ACMTF corresponds to $\pi=1$.

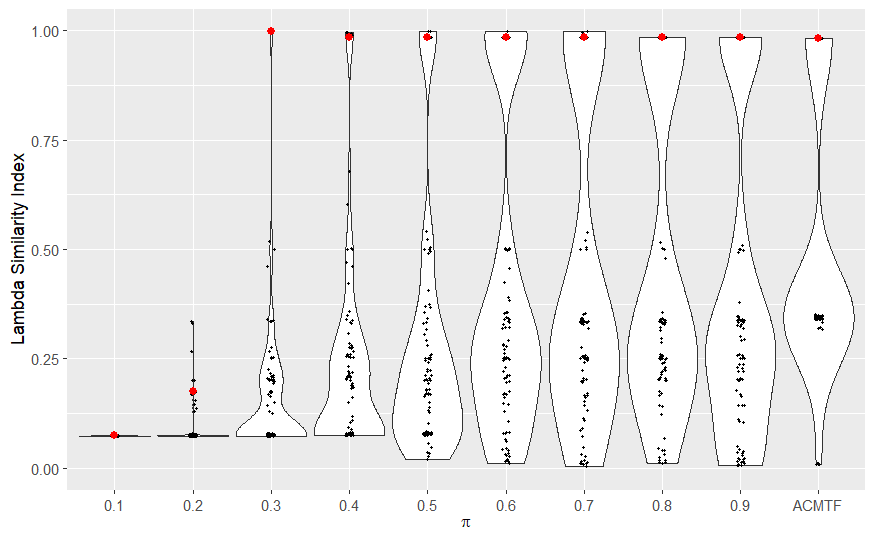

**Supplementary Figure 7.** Violin plots showing the Lambda Similarity Index observed across the randomly initialised ACMTF and ACMTF-R models in the first simulation. The model with the lowest loss per setting of $\pi$ is indicated with a red data point.

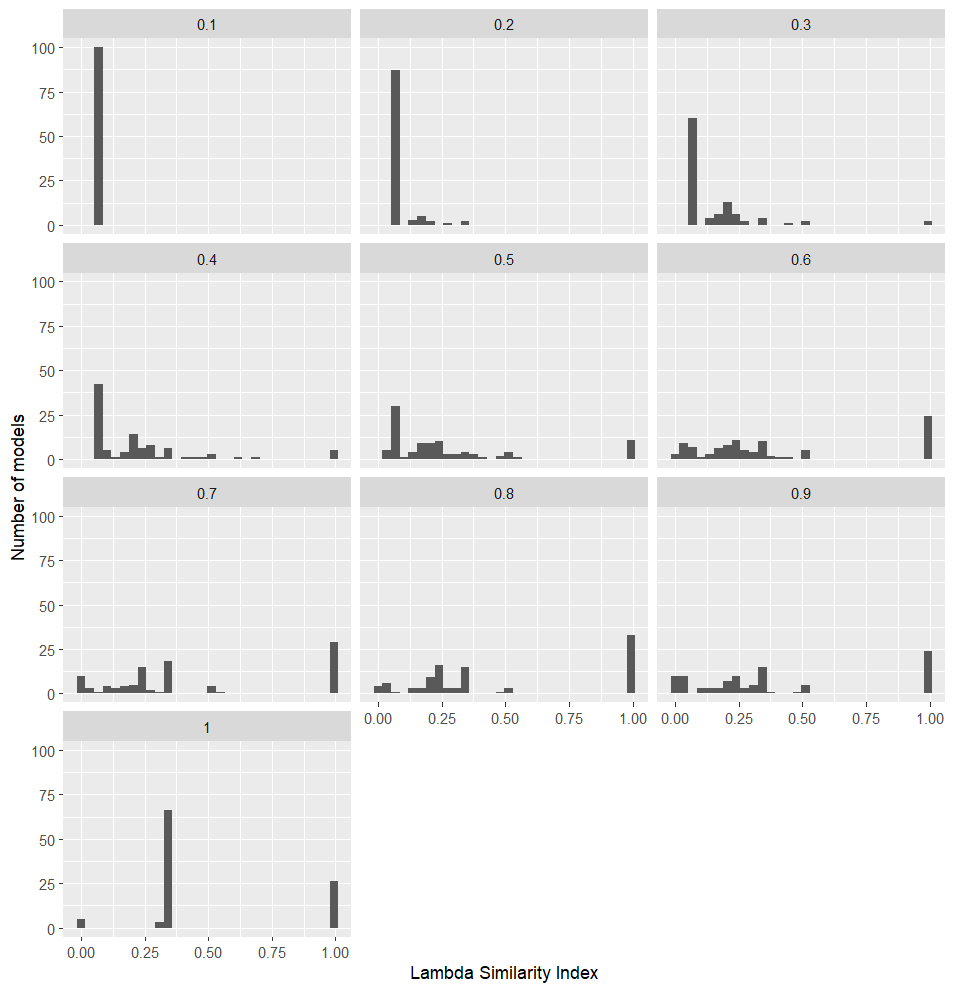

**Supplementary Figure 8.** Histogram of the $LSI$ observed across the randomly initialised ACMTF and ACMTF-R models in the first simulation. Plots are faceted by $\pi$ value and each contain 100 randomly initialised models. ACMTF corresponds to $\pi=1$.

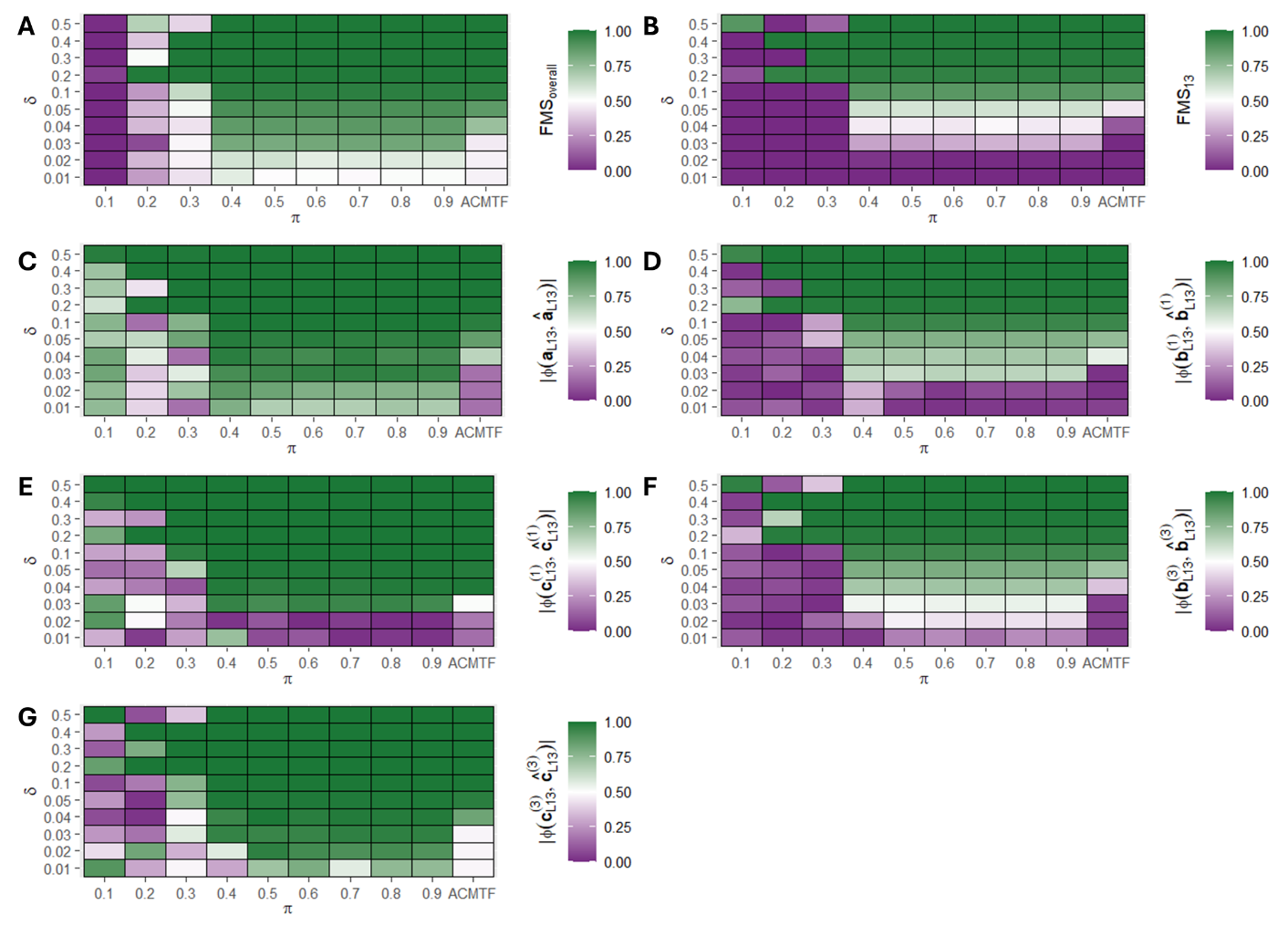

**Supplementary Figure 9. Factor recovery for varying sizes of the hidden component** $\boldsymbol{L}_{\boldsymbol{13}}$ **in ACMTF and ACMTF-R.** One hundred randomly initialised eight-component ACMTF-R and ACMTF models were fitted for each combination of hidden component size ($\delta$) and tuning ($\pi$), after which the lowest-loss models were compared. Outcome per setting for (A) overall factor recovery, (B) factor recovery of the hidden component, (C) shared subject mode recovery of the hidden component, (D) feature mode recovery of ${\underline{\boldsymbol{X}}}^{(1)}$ for the hidden component, (E) time mode recovery of ${\underline{\boldsymbol{X}}}^{(1)}$ for the hidden component, (F) feature mode recovery of ${\underline{\boldsymbol{X}}}^{(3)}$ for the hidden component, (G) time mode recovery of ${\underline{\boldsymbol{X}}}^{(3)}$ for the hidden component.

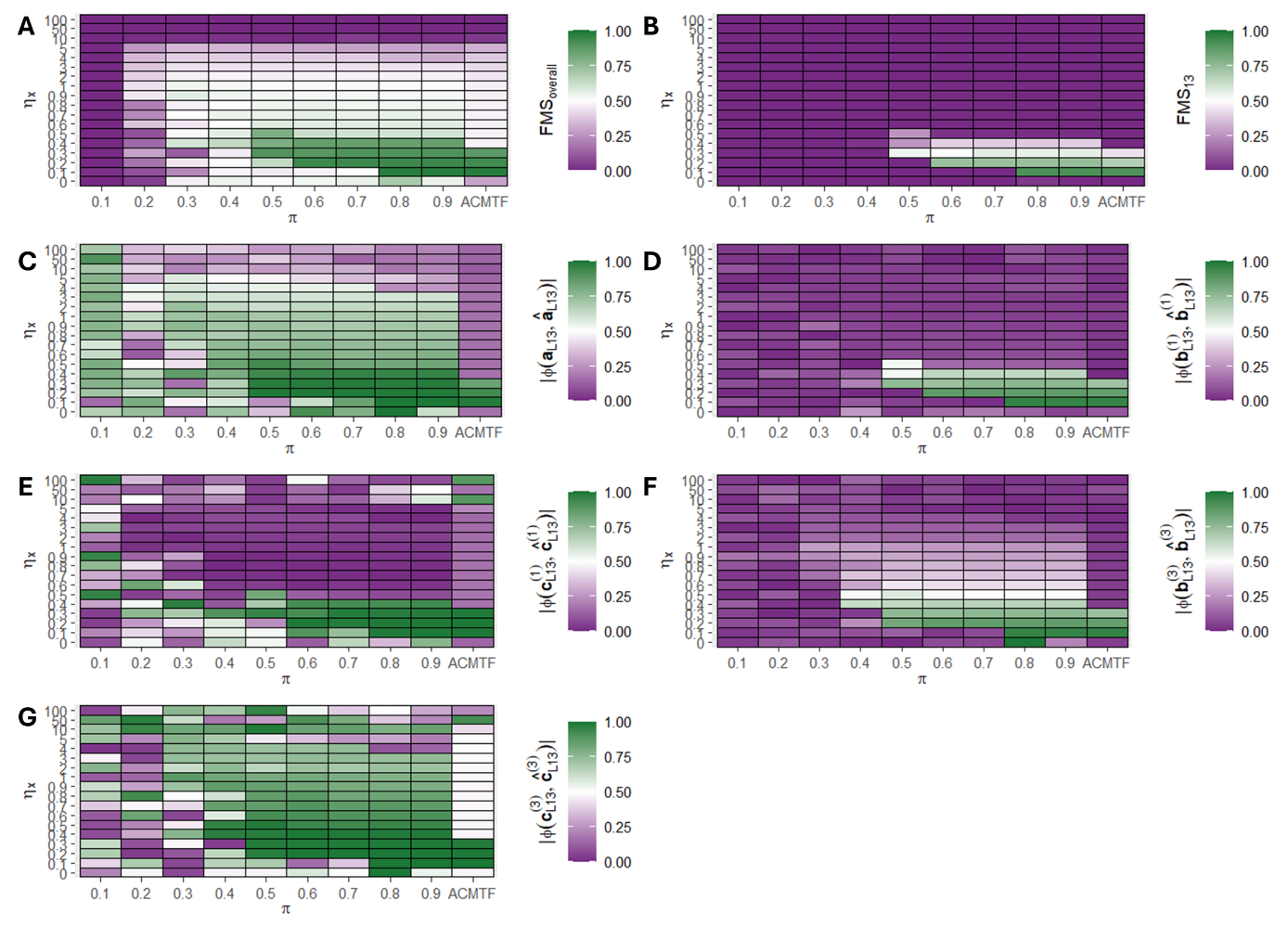

**Supplementary Figure 10. Factor recovery in ACMTF and ACMTF-R for varying noise levels on X.** One hundred randomly initialised eight-component ACMTF-R and ACMTF models were fitted for each combination of noise on X ($\eta_{X}$) and tuning parameter ($\pi$), after which the lowest-loss models were compared. Outcome per setting for (A) overall factor recovery, (B) factor recovery of the hidden component, (C) shared subject mode recovery of the hidden component, (D) feature mode recovery of ${\underline{\boldsymbol{X}}}^{(1)}$ for the hidden component, (E) time mode recovery of ${\underline{\boldsymbol{X}}}^{(1)}$ for the hidden component, (F) feature mode recovery of ${\underline{\boldsymbol{X}}}^{(3)}$ for the hidden component, (G) time mode recovery of ${\underline{\boldsymbol{X}}}^{(3)}$ for the hidden component.

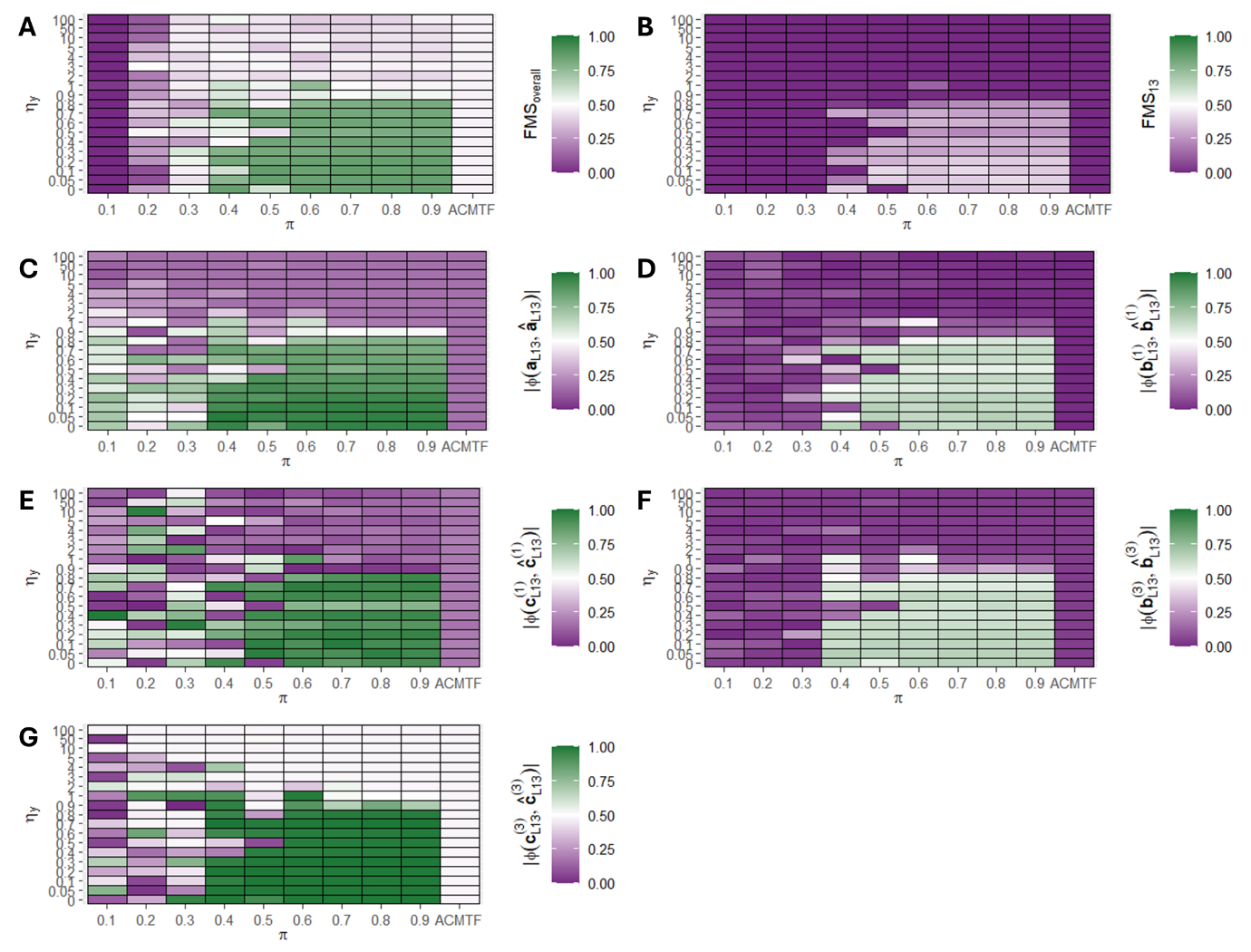

**Supplementary Figure 11. Factor recovery in ACMTF and ACMTF-R for varying noise levels on** $\boldsymbol{y}$**.** One hundred randomly initialised eight-component ACMTF-R and ACMTF models were fitted for each combination of noise on $\boldsymbol{y}$ ($\eta_{y}$) and tuning parameter ($\pi$), after which the lowest-loss models were compared. Outcome per setting for (A) overall factor recovery, (B) factor recovery of the hidden component, (C) shared subject mode recovery of the hidden component, (D) feature mode recovery of ${\underline{\boldsymbol{X}}}^{(1)}$ for the hidden component, (E) time mode recovery of ${\underline{\boldsymbol{X}}}^{(1)}$ for the hidden component, (F) feature mode recovery of ${\underline{\boldsymbol{X}}}^{(3)}$ for the hidden component, (G) time mode recovery of ${\underline{\boldsymbol{X}}}^{(3)}$ for the hidden component.

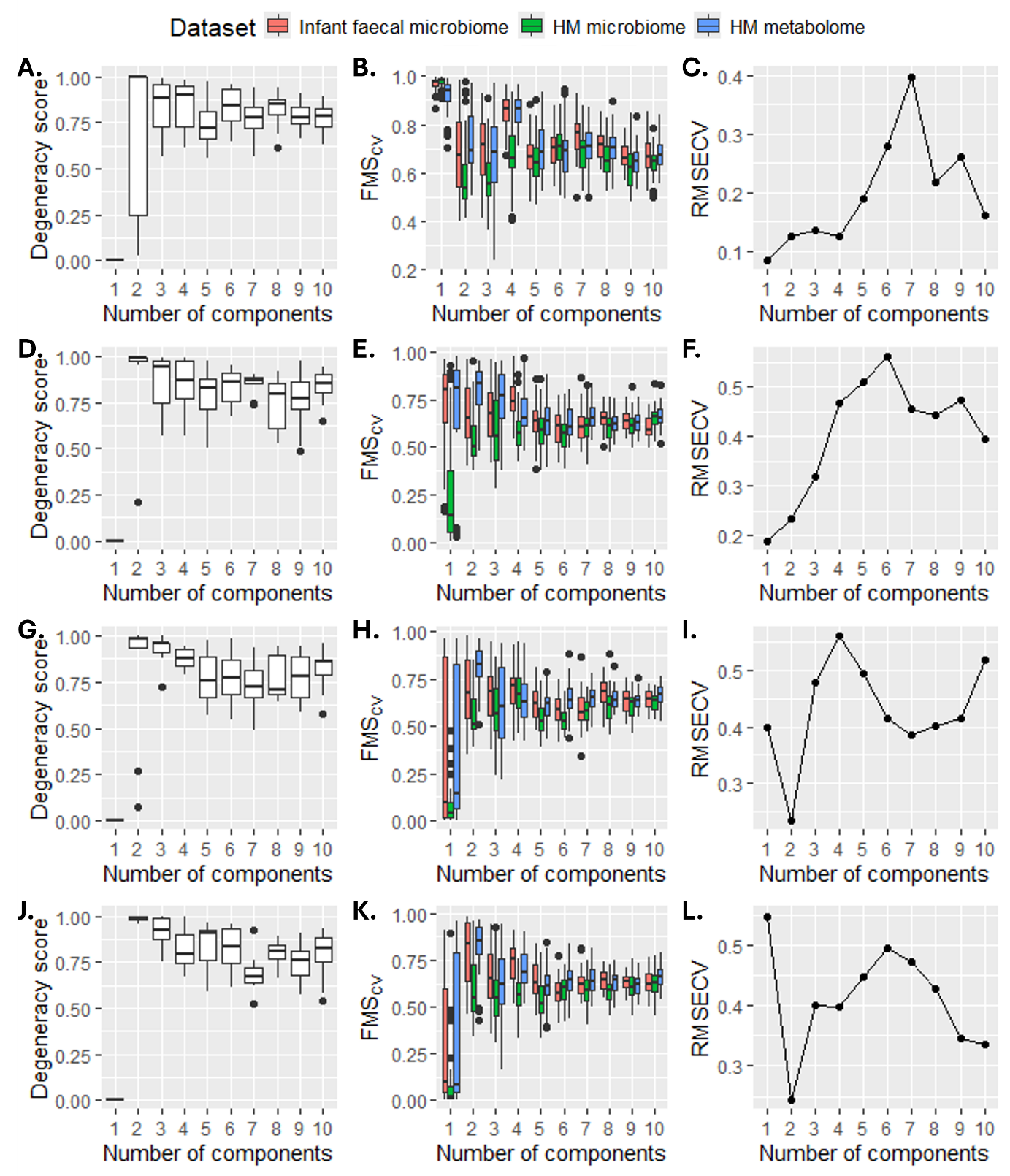

**Supplementary Figure 12.** Model selection to determine the correct number of components for ACMTF-R regressing on maternal ppBMI at (A-C) $\pi=0.80$, (D-F) $\pi=0.85$, (G-I) $\pi=0.90$, (J-L) $\pi=0.95$.

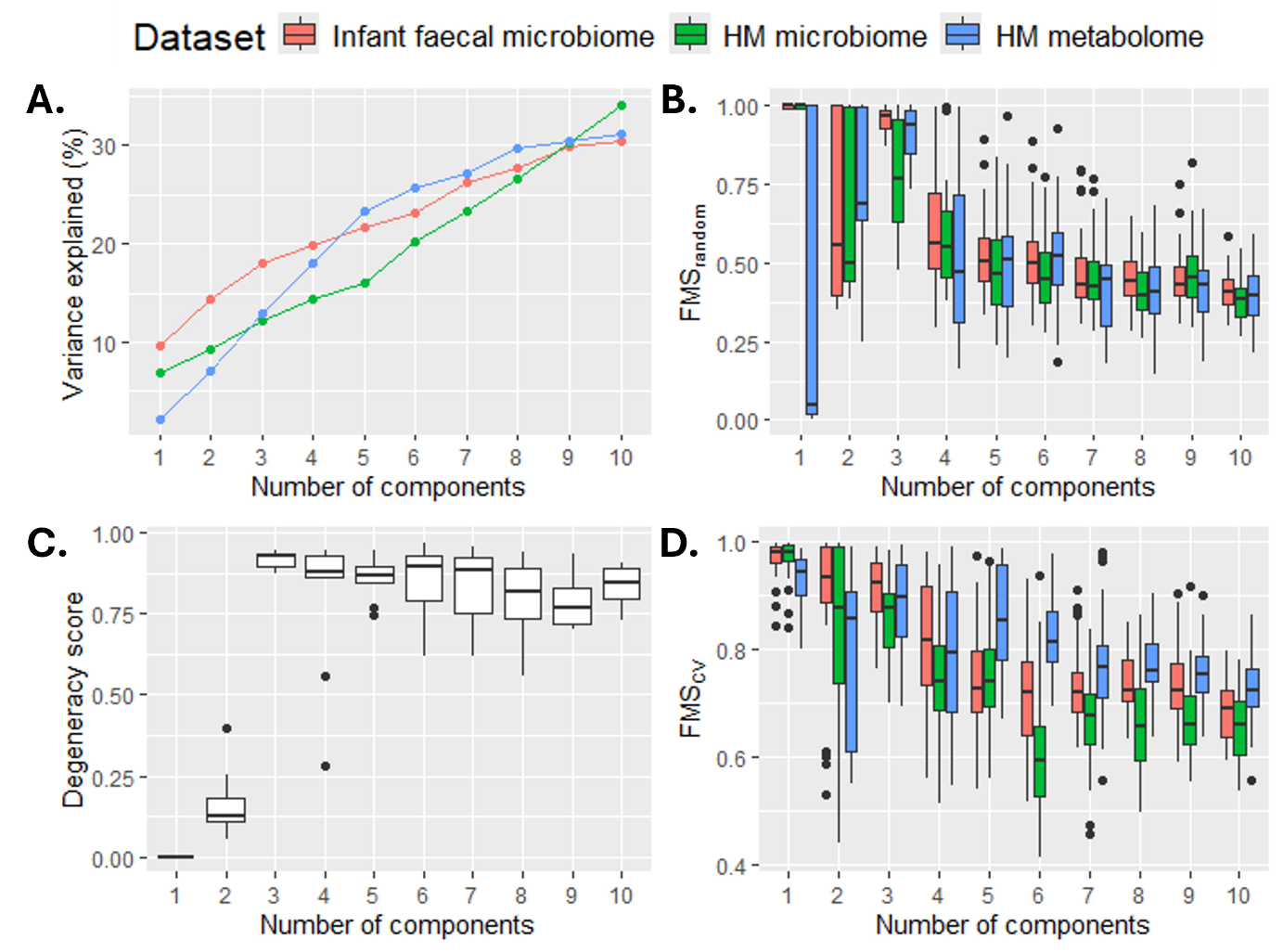

**Supplementary Figure 13.** Model selection outcome for the biological application using ACMTF to determine the appropriate number of components. (A) Variance explained per block. (B) $FMS_{random}$ across 10 randomly initialised models per setting. (C) Degeneracy score across the (same) randomly initialised models. (D) $FMS_{CV}$ across 10 CV folds.

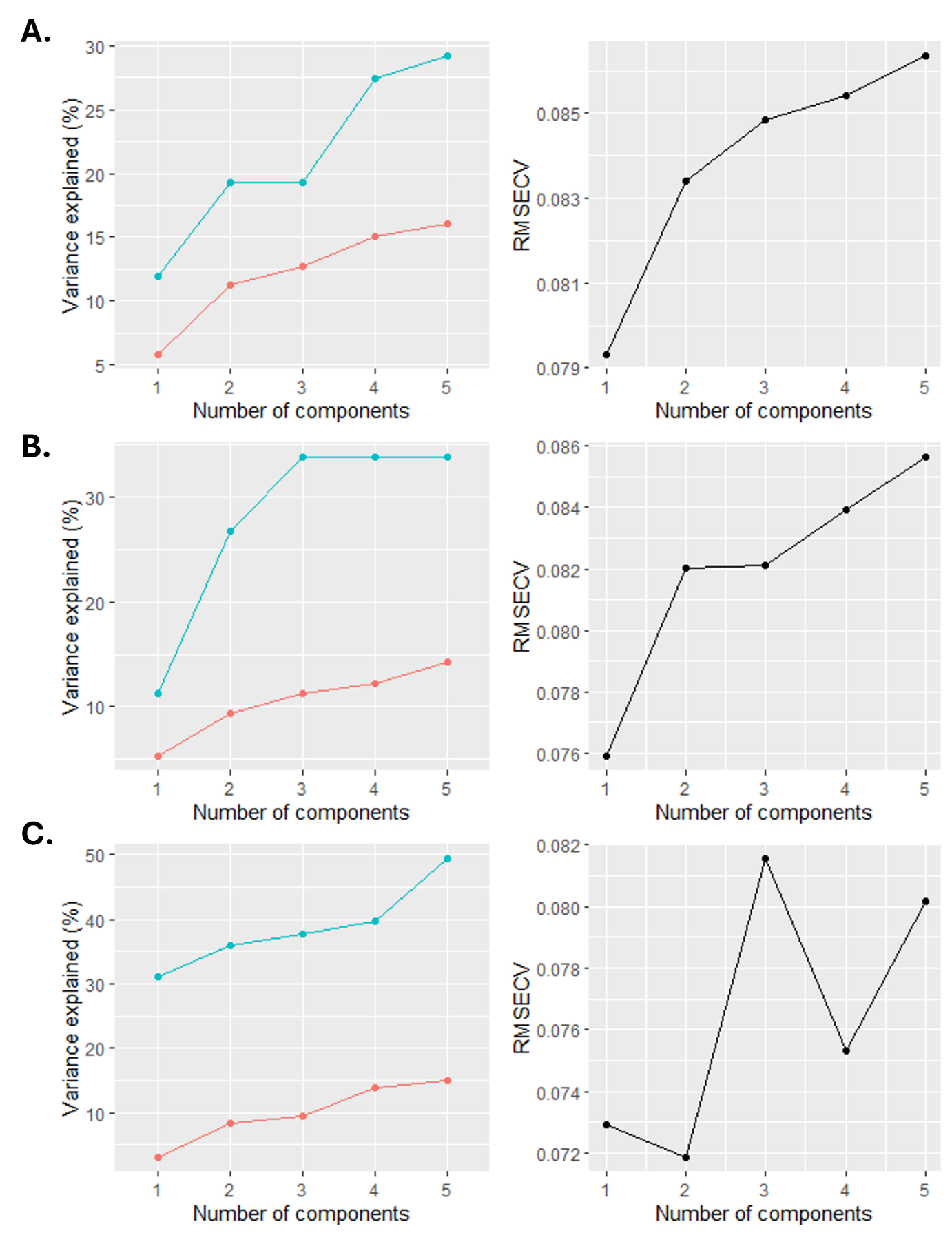

**Supplementary Figure 14**. Cross-validation outcome for NPLS for the (A) infant faecal microbiome, (B) HM milk, (C) HM metabolomics datasets of the biological application.

**Supplementary Tables**

**Supplementary Table 1.** Results of a Kendall’s tau test for the feature mode loadings of the lowest-loss ACMTF-R models against the true feature mode loadings per data block for the hidden local component $L_{13}$. Values correspond to $\tau$ and stars indicate their significance (*: $p<0.05$, **: $p<0.01$, ***: $p<0.001$).

| $\pi$ | ${\underline{\boldsymbol{X}}}^{(1)}$ | | ${\underline{\boldsymbol{X}}}^{(3)}$ | |
| --- | --- | --- | --- | --- |
|  | $\tau$ | 95% CI | $\tau$ | 95% CI |
| 1.00 (ACMTF) | 0.00 | -0.10 – 0.09 | -0.04 | -0.13 – 0.06 |
| 0.90 | 0.44*** | 0.37 – 0.51 | 0.36*** | 0.27 – 0.45 |
| 0.80 | 0.45*** | 0.38 – 0.51 | 0.36*** | 0.27 – 0.45 |
| 0.70 | 0.45*** | 0.38 – 0.52 | 0.37*** | 0.28 – 0.45 |
| 0.60 | 0.45*** | 0.38 – 0.52 | 0.36*** | 0.28 – 0.44 |
| 0.50 | 0.43*** | 0.35 – 0.50 | 0.35*** | 0.26 – 0.43 |
| 0.40 | 0.43*** | 0.35 – 0.50 | 0.35*** | 0.26 – 0.44 |
| 0.30 | 0.03 | 0.08 – 0.12 | 0.00 | -0.10 – 0.10 |
| 0.20 | 0.09 | 0.00 – 0.18 | -0.05 | -0.15 – 0.04 |
| 0.10 | 0.02 | -0.08 – 0.11 | -0.06 | -0.15 – 0.03 |

**Supplementary Table 2**. Correlation matrix of the MAINHEALTH metadata. Birth mode, secretor status (*Se*), and Lewis status (*Le*) are binarized variables.

|  | ppBMI | Birth mode | *Se* | *Le* | WHZ |
| --- | --- | --- | --- | --- | --- |
| ppBMI | 1 | 0.04 | -0.02 | 0.06 | 0.07 |
| Birth mode |  | 1 | -0.06 | -0.05 | 0.03 |
| *Se* |  |  | 1 | -0.10 | -0.08 |
| *Le* |  |  |  | 1 | -0.18 |
| WHZ |  |  |  |  | 1 |
