## Appendix 1 for "ACMTF-R: supervised multi-omics data integration uncovering shared and distinct outcome-associated variation"

### ACMTF-R gradient derivation

G.R. van der Ploeg

March 2025

#### 1 Introduction

In this text, the derivative of the ACMTF-R loss function is reported. This is done by first repeating the loss function of ACMTF-R and subsequently writing out some of the terms for convenience. Then the gradient is derived term by term.

The loss function of ACMTF-R is repeated here for convenience to the reader, where the assumptions that every term is multiplied by $1/2$ and that the first mode loadings $\boldsymbol{A}$ are shared between all blocks are applied. In addition, each of the six terms are defined as separate functions $f_1, f_2, ..., f_6$ such that later derivations can be done per term.

$$
\begin{aligned}
f(\alpha,\beta,\epsilon,\pi,\boldsymbol{\Lambda},\boldsymbol{\rho},\boldsymbol{A},\boldsymbol{B}^{(1)},\boldsymbol{C}^{(1)},...,\boldsymbol{B}^{(P)},\boldsymbol{C}^{(P)}) = & \underbrace{\frac{1}{2}\pi\sum_{p=1}^{P}\|\underline{\boldsymbol{X}}^{(p)}-\hat{\underline{\boldsymbol{X}}}^{(p)}\|^2}_{f_1} \\
& \underbrace{+\frac{1}{2}(1-\pi)\|\boldsymbol{y}-\boldsymbol{A}\boldsymbol{\rho}\|^2}_{f_2} \\
& \underbrace{+\frac{1}{2}\beta\sum_{p=1}^{P}\sum_{f=1}^{F}\sqrt{\lambda_{fp}^2+\epsilon}}_{f_3} \\
& \underbrace{+\frac{1}{2}\alpha\sum_{p=1}^{P}\sum_{f=1}^{F}(\|\boldsymbol{a}_f\|-1)^2}_{f_4} \\
& \underbrace{+\frac{1}{2}\alpha\sum_{p=1}^{P}\sum_{f=1}^{F}(\|\boldsymbol{b}_f^{(p)}\|-1)^2}_{f_5} \\
& \underbrace{+\frac{1}{2}\alpha\sum_{p=1}^{P}\sum_{f=1}^{F}(\|\boldsymbol{c}_f^{(p)}\|-1)^2}_{f_6}
\end{aligned} \tag{1}
$$

where the $F \times P$ matrix $\boldsymbol{\Lambda} = [\boldsymbol{\lambda}_1\ \boldsymbol{\lambda}_2\ \cdots\ \boldsymbol{\lambda}_p]$ specifies the norms per component and per data block, $\alpha > 0$ is a penalty setting for norm-1 components, $\beta > 0$ is a sparsity penalty setting for $\boldsymbol{\Lambda}$, $\epsilon > 0$ is a small term added to $\boldsymbol{\Lambda}$ elements to make the loss function differentiable, $\pi$ is the tuning parameter, $\boldsymbol{\rho}$ is the vector of the regression coefficients of the shared subject loadings in $\boldsymbol{A}$ onto $\boldsymbol{y}$, $F$ is the number of components, and $P$ is the number of blocks.

The aim of this section is to compute the gradient of the loss function

$$
\begin{aligned}
\nabla f = [\text{vec}(\frac{\partial f}{\partial \boldsymbol{A}})^T\ \text{vec}(\frac{\partial f}{\partial \boldsymbol{B}^{(1)}})^T\ \text{vec}(\frac{\partial f}{\partial \boldsymbol{C}^{(1)}})^T\ \cdots \text{vec}(\frac{\partial f}{\partial \boldsymbol{B}^{(P)}})^T\ \text{vec}(\frac{\partial f}{\partial \boldsymbol{C}^{(P)}})^T \\
(\frac{\partial f}{\partial \boldsymbol{\lambda}_1})^T\ (\frac{\partial f}{\partial \boldsymbol{\lambda}_2})^T\ \cdots\ (\frac{\partial f}{\partial \boldsymbol{\lambda}_P})^T]^T.
\end{aligned} \tag{2}
$$

which will be done by computing this derivative per term.

We briefly note here that the partial derivative of $f$ with respect to $\boldsymbol{\lambda}_1, \boldsymbol{\lambda}_2, ..., \boldsymbol{\lambda}_p$ is the same as originally derived for ACMTF [1] and will not be repeated here.

#### 2 Useful equations

The first term of the loss function $f_1$ can be expressed element-wise as

$$
\begin{aligned}
f_1 &= \frac{1}{2}\pi \sum_{p=1}^{P} \|\underline{\boldsymbol{X}}^{(p)} - \hat{\underline{\boldsymbol{X}}}^{(p)}\|^2 \\
&= \frac{1}{2}\pi \sum_{p=1}^{P} \left( \sqrt{\sum_{i=1}^{I}\sum_{j=1}^{J}\sum_{k=1}^{K} (x_{ijk}^{(p)} - \hat{x}_{ijk}^{(p)})^2} \right)^2 \\
&= \frac{1}{2}\pi \sum_{p=1}^{P}\sum_{i=1}^{I}\sum_{j=1}^{J}\sum_{k=1}^{K} (x_{ijk}^{(p)} - \hat{x}_{ijk}^{(p)})^2 \\
&= \frac{1}{2}\pi \sum_{p=1}^{P}\sum_{i=1}^{I}\sum_{j=1}^{J}\sum_{k=1}^{K} (x_{ijk}^{(p)} - \sum_{f=1}^{F} \lambda_{f,p} a_{if} b_{jf}^{(p)} c_{kf}^{(p)})^2 \\
&= \frac{1}{2}\,\pi \sum_{p=1}^{P}\sum_{i=1}^{I}\sum_{j=1}^{J}\sum_{k=1}^{K} \Bigg[ (x_{ijk}^{(p)})^2 - 2x_{ijk}^{(p)} \sum_{f=1}^{F} \lambda_{f,p} a_{if} b_{jf}^{(p)} c_{kf}^{(p)} \\
&+ (\sum_{f=1}^{F} \lambda_{f,p} a_{if} b_{jf}^{(p)} c_{kf}^{(p)})^2 \Bigg] .
\end{aligned}
\tag{3}
$$

The second term of the loss function $f_2$ can be expressed element-wise as

$$
\begin{aligned}
f_2 &= \frac{1}{2}(1-\pi)\|\boldsymbol{y} - \boldsymbol{A\rho}\|^2 \\
&= \frac{1}{2}(1-\pi)\sqrt{\sum_{i=1}^{I}\left( y_i - \sum_{f=1}^{F} a_{if}\rho_f \right)^2}^2 \\
&= \frac{1}{2}(1-\pi)\sum_{i=1}^{I}\left( y_i - \sum_{f=1}^{F} a_{if}\rho_f \right)^2 \\
&= \frac{1}{2}(1-\pi)\sum_{i=1}^{I}\left[ y_i^2 - 2y_i \sum_{f=1}^{F} a_{if}\rho_f + (\sum_{f=1}^{F} a_{if}\rho_f)^2 \right] .
\end{aligned}
\tag{4}
$$

#### 3 Derivative of A

In this section the partial derivative of the loss function with respect to $\boldsymbol{A}$ is reported through

$$
\begin{aligned}
\frac{\partial f}{\partial \boldsymbol{A}} &= \frac{\partial f_1}{\partial \boldsymbol{A}} + \frac{\partial f_2}{\partial \boldsymbol{A}} + \frac{\partial f_3}{\partial \boldsymbol{A}} + \frac{\partial f_4}{\partial \boldsymbol{A}} + \frac{\partial f_5}{\partial \boldsymbol{A}} + \frac{\partial f_6}{\partial \boldsymbol{A}} \\
&= \frac{\partial f_1}{\partial \boldsymbol{A}} + \frac{\partial f_2}{\partial \boldsymbol{A}} + \frac{\partial f_4}{\partial \boldsymbol{A}}
\end{aligned} \tag{5}
$$

where $\frac{\partial f_3}{\partial \boldsymbol{A}} = \frac{\partial f_5}{\partial \boldsymbol{A}} = \frac{\partial f_6}{\partial \boldsymbol{A}} = 0$ since those terms of the loss function do not contain relationships with $\boldsymbol{A}$.

The element-wise definition of $f_1$ in Equation 3 can be used to express the partial derivative with respect to $\boldsymbol{A}$ as

$$
\begin{aligned}
\frac{\partial f_1}{\partial a_{if}} &= \frac{1}{2}\pi \sum_{i=1}^{I}\sum_{j=1}^{J}\sum_{k=1}^{K}\left[-2x_{ijk}^{(p)} \sum_{f=1}^{F} \lambda_{f,p} b_{jf}^{(p)} c_{kf}^{(p)}\right. \\
&\quad \left. +2(\sum_{f=1}^{F} \lambda_{f,p} a_{if} b_{jf}^{(p)} c_{kf}^{(p)})(\sum_{f=1}^{F} \lambda_{f,p} b_{jf}^{(p)} c_{kf}^{(p)})\right] \\
&= \pi \sum_{i=1}^{I}\sum_{j=1}^{J}\sum_{k=1}^{K}\left[(-x_{ijk}^{(p)} + \sum_{f=1}^{F} \lambda_{f,p} a_{if} b_{jf}^{(p)} c_{kf}^{(p)})(\sum_{f=1}^{F} \lambda_{f,p} b_{jf}^{(p)} c_{kf}^{(p)})\right] \\
&= \pi \sum_{i=1}^{I}\sum_{j=1}^{J}\sum_{k=1}^{K}\left[(\sum_{f=1}^{F} \lambda_{f,p} a_{if} b_{jf}^{(p)} c_{kf}^{(p)} - x_{ijk}^{(p)})(\sum_{f=1}^{F} \lambda_{f,p} b_{jf}^{(p)} c_{kf}^{(p)})\right]
\end{aligned} \tag{6}
$$

which can then be written in matrix notation as

$$
\frac{\partial f_1}{\partial \boldsymbol{A}} = \pi \sum_{p=1}^{P} (\hat{\boldsymbol{X}}_a^{(p)} - \boldsymbol{X}_a^{(p)})(\boldsymbol{\lambda}_p \odot \boldsymbol{C}^{(p)} \odot \boldsymbol{B}^{(p)}). \tag{7}
$$

The element-wise definition of $f_2$ in Equation 4 can be used to express the partial derivative with respect to $\boldsymbol{A}$ as

$$
\begin{aligned}
\frac{\partial f_2}{\partial a_{if}} &= \frac{1}{2}(1-\pi) \sum_{i=1}^{I}\left[-2y_i \sum_{f=1}^{F} \rho_f + 2(\sum_{f=1}^{F} a_{if}\rho_f)(\sum_{f=1}^{F} \rho_f)\right] \\
&= (1-\pi) \sum_{i=1}^{I}\left[-y_i \sum_{f=1}^{F} \rho_f + (\sum_{f=1}^{F} a_{if}\rho_f)(\sum_{f=1}^{F} \rho_f)\right] \\
&= (1-\pi) \sum_{i=1}^{I}\left[(-y_i + \sum_{f=1}^{F} a_{if}\rho_f)(\sum_{f=1}^{F} \rho_f)\right] \\
&= (1-\pi) \sum_{i=1}^{I}\left[(\sum_{f=1}^{F} a_{if}\rho_f - y_i)(\sum_{f=1}^{F} \rho_f)\right]
\end{aligned} \tag{8}
$$

which can then be written in matrix notation as

$$\frac{\partial f_2}{\partial \boldsymbol{A}} = (1-\pi)(\boldsymbol{A}\boldsymbol{\rho} - \boldsymbol{y})\boldsymbol{\rho}^T. \tag{9}$$

The partial derivative of $f_4$ with respect to $\boldsymbol{A}$ is the same as originally defined [1] for ACMTF

$$\frac{\partial f_4}{\partial \boldsymbol{A}} = \alpha(\boldsymbol{A} - \bar{\boldsymbol{A}}). \tag{10}$$

Combining the terms as defined in Equations 7, 9, and 10, this yields the full partial derivative of the loss function $f$ with respect to $\boldsymbol{A}$

$$\begin{aligned}\frac{\partial f}{\partial \boldsymbol{A}} &= \frac{\partial f_1}{\partial \boldsymbol{A}} + \frac{\partial f_2}{\partial \boldsymbol{A}} + \frac{\partial f_4}{\partial \boldsymbol{A}} \\ &= \pi \sum_{p=1}^{P} (\hat{\boldsymbol{X}}_a^{(p)} - \boldsymbol{X}_a^{(p)})(\boldsymbol{\lambda}_p \odot \boldsymbol{C}^{(p)} \odot \boldsymbol{B}^{(p)}) \\ &+ (1-\pi)(\boldsymbol{A}\boldsymbol{\rho} - \boldsymbol{y})\boldsymbol{\rho}^T + \alpha(\boldsymbol{A} - \bar{\boldsymbol{A}}).\end{aligned} \tag{11}$$

### 4 Derivative of B

In this section the partial derivative of the loss function with respect to $\boldsymbol{B}^{(p)}$ is reported through

$$\begin{aligned}\frac{\partial f}{\partial \boldsymbol{B}^{(p)}} &= \frac{\partial f_1}{\partial \boldsymbol{B}^{(p)}} + \frac{\partial f_2}{\partial \boldsymbol{B}^{(p)}} + \frac{\partial f_3}{\partial \boldsymbol{B}^{(p)}} + \frac{\partial f_4}{\partial \boldsymbol{B}^{(p)}} + \frac{\partial f_5}{\partial \boldsymbol{B}^{(p)}} + \frac{\partial f_6}{\partial \boldsymbol{B}^{(p)}} \\ &= \frac{\partial f_1}{\partial \boldsymbol{B}^{(p)}} + \frac{\partial f_5}{\partial \boldsymbol{B}^{(p)}}\end{aligned} \tag{12}$$

where $\frac{\partial f_2}{\partial \boldsymbol{B}^{(p)}} = \frac{\partial f_3}{\partial \boldsymbol{B}^{(p)}} = \frac{\partial f_4}{\partial \boldsymbol{B}^{(p)}} = \frac{\partial f_6}{\partial \boldsymbol{B}^{(p)}} = 0$ since those terms of the loss function do not contain relationships with $\boldsymbol{B}^{(p)}$.

The element-wise definition of $f_1$ in Equation 3 can be used to express the partial derivative with respect to $\boldsymbol{B}^{(p)}$ as

$$
\begin{aligned}
\frac{\partial f_1}{\partial b_{jf}^{(p)}} &= \frac{1}{2}\pi \sum_{i=1}^{I}\sum_{j=1}^{J}\sum_{k=1}^{K} \left[ -2x_{ijk}^{(p)} \sum_{f=1}^{F} \lambda_{f,p} a_{if} c_{kf}^{(p)} \right. \\
&\left. +2(\sum_{f=1}^{F} \lambda_{f,p} a_{if} b_{jf}^{(p)} c_{kf}^{(p)})(\sum_{f=1}^{F} \lambda_{f,p} a_{if} c_{kf}^{(p)}) \right] \\
&= \pi \sum_{i=1}^{I}\sum_{j=1}^{J}\sum_{k=1}^{K} \left[ (-x_{ijk}^{(p)} + \sum_{f=1}^{F} \lambda_{f,p} a_{if} b_{jf}^{(p)} c_{kf}^{(p)})(\sum_{f=1}^{F} \lambda_{f,p} a_{if} c_{kf}^{(p)}) \right] \\
&= \pi \sum_{i=1}^{I}\sum_{j=1}^{J}\sum_{k=1}^{K} \left[ (\sum_{f=1}^{F} \lambda_{f,p} a_{if} b_{jf}^{(p)} c_{kf}^{(p)} - x_{ijk}^{(p)})(\sum_{f=1}^{F} \lambda_{f,p} a_{if} c_{kf}^{(p)}) \right]
\end{aligned} \tag{13}
$$

which can then be written in matrix notation as

$$
\frac{\partial f_1}{\partial \boldsymbol{B}^{(p)}} = \pi \sum_{p=1}^{P} (\hat{\boldsymbol{X}}_a^{(p)} - \boldsymbol{X}_a^{(p)})(\boldsymbol{\lambda}_p \odot \boldsymbol{C}^{(p)} \odot \boldsymbol{A}). \tag{14}
$$

The partial derivative of $f_5$ with respect to $\boldsymbol{B}^{(p)}$ is the same as originally defined [1] for ACMTF

$$
\frac{\partial f_5}{\partial \boldsymbol{B}^{(p)}} = \alpha(\boldsymbol{B}^{(p)} - \bar{\boldsymbol{B}}^{(p)}). \tag{15}
$$

Combining the terms as defined in Equations 14 and 15, this yields the full partial derivative of the loss function $f$ with respect to $\boldsymbol{B}^{(p)}$

$$
\begin{aligned}
\frac{\partial f}{\partial \boldsymbol{B}^{(p)}} &= \frac{\partial f_1}{\partial \boldsymbol{B}^{(p)}} + \frac{\partial f_5}{\partial \boldsymbol{B}^{(p)}} \\
&= \pi \sum_{p=1}^{P} (\hat{\boldsymbol{X}}_a^{(p)} - \boldsymbol{X}_a^{(p)})(\boldsymbol{\lambda}_p \odot \boldsymbol{C}^{(p)} \odot \boldsymbol{A}) + \alpha(\boldsymbol{B}^{(p)} - \bar{\boldsymbol{B}}^{(p)}).
\end{aligned} \tag{16}
$$

### 5 Derivative of C

In this section the partial derivative of the loss function with respect to $\boldsymbol{C}^{(p)}$ is reported through

$$
\begin{aligned}
\frac{\partial f}{\partial \boldsymbol{C}^{(p)}} &= \frac{\partial f_1}{\partial \boldsymbol{C}^{(p)}} + \frac{\partial f_2}{\partial \boldsymbol{C}^{(p)}} + \frac{\partial f_3}{\partial \boldsymbol{C}^{(p)}} + \frac{\partial f_4}{\partial \boldsymbol{C}^{(p)}} + \frac{\partial f_5}{\partial \boldsymbol{C}^{(p)}} + \frac{\partial f_6}{\partial \boldsymbol{C}^{(p)}} \\
&= \frac{\partial f_1}{\partial \boldsymbol{C}^{(p)}} + \frac{\partial f_6}{\partial \boldsymbol{C}^{(p)}}
\end{aligned} \tag{17}
$$

where $\frac{\partial f_2}{\partial \boldsymbol{C}^{(p)}} = \frac{\partial f_3}{\partial \boldsymbol{C}^{(p)}} = \frac{\partial f_4}{\partial \boldsymbol{C}^{(p)}} = \frac{\partial f_5}{\partial \boldsymbol{C}^{(p)}} = 0$ since those terms of the loss function do not contain relationships with $\boldsymbol{C}^{(p)}$.

The element-wise definition of $f_1$ in Equation 3 can be used to express the partial derivative with respect to $\boldsymbol{C}^{(p)}$ as

$$\begin{aligned}\frac{\partial f_1}{\partial c_{kf}^{(p)}} &= \frac{1}{2}\pi \sum_{i=1}^{I}\sum_{j=1}^{J}\sum_{k=1}^{K}\Bigg[-2x_{ijk}^{(p)} \sum_{f=1}^{F} \lambda_{f,p} b_{jf}^{(p)} c_{kf}^{(p)} \\ &\quad +2(\sum_{f=1}^{F} \lambda_{f,p} a_{if} b_{jf}^{(p)} c_{kf}^{(p)})(\sum_{f=1}^{F} \lambda_{f,p} b_{jf}^{(p)} c_{kf}^{(p)})\Bigg] \\ &= \pi \sum_{i=1}^{I}\sum_{j=1}^{J}\sum_{k=1}^{K}\Bigg[(-x_{ijk}^{(p)} + \sum_{f=1}^{F} \lambda_{f,p} a_{if} b_{jf}^{(p)} c_{kf}^{(p)})(\sum_{f=1}^{F} \lambda_{f,p} b_{jf}^{(p)} c_{kf}^{(p)})\Bigg] \\ &= \pi \sum_{i=1}^{I}\sum_{j=1}^{J}\sum_{k=1}^{K}\Bigg[(\sum_{f=1}^{F} \lambda_{f,p} a_{if} b_{jf}^{(p)} c_{kf}^{(p)} - x_{ijk}^{(p)})(\sum_{f=1}^{F} \lambda_{f,p} b_{jf}^{(p)} c_{kf}^{(p)})\Bigg]\end{aligned} \tag{18}$$

which can then be written in matrix notation as

$$\frac{\partial f_1}{\partial \boldsymbol{C}^{(p)}} = \pi \sum_{p=1}^{P} (\hat{\boldsymbol{X}}_a^{(p)} - \boldsymbol{X}_a^{(p)})(\boldsymbol{\lambda}_p \odot \boldsymbol{C}^{(p)} \odot \boldsymbol{B}^{(p)}). \tag{19}$$

The partial derivative of $f_5$ with respect to $\boldsymbol{C}^{(p)}$ is the same as originally defined [1] for ACMTF

$$\frac{\partial f_6}{\partial \boldsymbol{C}^{(p)}} = \alpha(\boldsymbol{C}^{(p)} - \bar{\boldsymbol{C}}^{(p)}). \tag{20}$$

Combining the terms as defined in Equations 19 and 20, this yields the full partial derivative of the loss function $f$ with respect to $\boldsymbol{C}^{(p)}$

$$\begin{aligned}\frac{\partial f}{\partial \boldsymbol{C}^{(p)}} &= \frac{\partial f_1}{\partial \boldsymbol{C}^{(p)}} + \frac{\partial f_6}{\partial \boldsymbol{C}^{(p)}} \\ &= \pi \sum_{p=1}^{P} (\hat{\boldsymbol{X}}_a^{(p)} - \boldsymbol{X}_a^{(p)})(\boldsymbol{\lambda}_p \odot \boldsymbol{C}^{(p)} \odot \boldsymbol{B}^{(p)}) + \alpha(\boldsymbol{C}^{(p)} - \bar{\boldsymbol{C}}^{(p)}).\end{aligned} \tag{21}$$
